# 5-Aza-Cytidine Enhances Terminal Polyadenylation Site Usage for Full-Length Transcripts in Cells

**DOI:** 10.1101/2024.02.22.581641

**Authors:** Samuel Ogunsola, Ling Liu, Urmi Das, Jiuyong Xie

## Abstract

As an inhibitor of DNA methyltransferases (DNMTs) and an anti-cancer drug, 5-aza-cytidine (5-azaC)’s many effects on gene expression remains unclear. Here, we show that 5-azaC treatment of cultured GH_3_ pituitary tumour cells increases relative usage of genomic terminal exons (GTEs) across the transcriptome. This effect is largely achieved by shifting mRNA polyadenylation from proximal poly(A) sites to GTEs, which harbour a more optimal consensus motif of poly(A) signals. Consistent with this shift, 5-azaC upregulates the mRNA anti-termination factors Scaf4 and Scaf8 while downregulating the early termination enhancer E2f2. In MOLM-13 leukaemia cells, 5-azaC similarly promotes the production of full-length transcripts and regulates alternative polyadenylation factors, some of which in the same direction as observed in GH_3_ cells. Moreover, PCF11, a factor known to promote proximal poly(A) site usage, is upregulated in both cell lines, suggesting a homeostatic response by these cells to counteract transcript lengthening during 5-azaC treatment. Together, these findings uncover a previously unrecognized effect of 5-azaC on gene expression: directional promotion of terminal polyadenylation site usage, driving a transcriptome-wide switch from shortened to full-length mRNAs in tumour or cancer cells and consequently altering the alternative usage of multiple 3′ exons.

## Introduction

5-aza-cytidine (5-azaC), an FDA-approved chemotherapeutic agent that binds and inactivates DNA methyltransferases (DNMTs), inhibits DNA methylation [1]. In addition to its well-established effects on DNA, 5-azaC influences multiple aspects of RNA metabolism [1]. However, its impact on RNA processing, particularly at the exon level, remains poorly understood.

DNA methylation and gene expression exhibit a gene-region-specific relationship: methylation level of promoters, first exons and first introns, but not the rest of gene body, is negatively correlated with gene expression [2, 3]. Whether 5-azaC exerts similarly region-dependent effects on exonic DNA methylation and exon usage has not been explored. This question motivated the present study and led to both expected and unexpected findings.

Here, we analysed RNA-Seq and whole-genome bisulfite sequencing (WGBS) data from GH_3_ pituitary tumour cells treated with 5-azaC. We focused on differential methylation and usage of genomic first, internal, and terminal exons (GTEs, to distinguish them from transcript-defined alternative terminal exons). Remarkably, 5-azaC promoted widespread usage of multiple 3′ exons and terminal polyadenylation sites, accompanied by more frequent changes of DNA methylation of GTEs. In parallel, 5-azaC altered the expression of genes involved in transcript length regulation and pre-mRNA processing. Importantly, similar effects were observed in a leukaemia cell line. Together, these findings suggest that 5-azaC may counteract transcript shortening in cancer cells through a unique, directional effect on 3′ exon usage, polyadenylation, and exon-specific DNA methylation, thereby restoring full-length mRNAs, their encoded protein domains, and associated functions.

## Results

### 1. Differential effects of 5-azaC on the relative exon usage of the genomic first, internal and terminal exons (GTE) of genes

To analyze potential exon-rank-dependent effects by the DNA methylation inhibitor 5-azaC, we treated GH_3_ pituitary tumor cells with 50μM 5-azaC and compared them with untreated controls [4]. Our analysis focused on the RNA sequencing data of genomic first, internal, and terminal exons by using DEXSeq [5], where first and terminal exons were designated as E001 for the positive (+) and negative (-) chromosome strands, respectively. For internal exons, we selected E004, E005, and E006 from the DEXSeq exon/bin list of genes with more than 8 exons. To further validate exon ranks, we randomly selected 100 genes from each group to verify in the Integrated Genomics Viewer [6]. This approach revealed 1709 first exons, 1790 internal exons, and 1874 terminal exons with a statistically significant difference upon 5-azaC treatment (Fig. 1A, *padj* < 0.05). Among the three groups of exons emerged an intriguing pattern: 5% of the first, 41% of the internal, and 92% of the terminal exons showed increased relative usage. This increasing percentages suggest a distinct exon-rank-dependent impact of 5-azaC on exon usage (Fig. 1A).

**Fig. 1.**
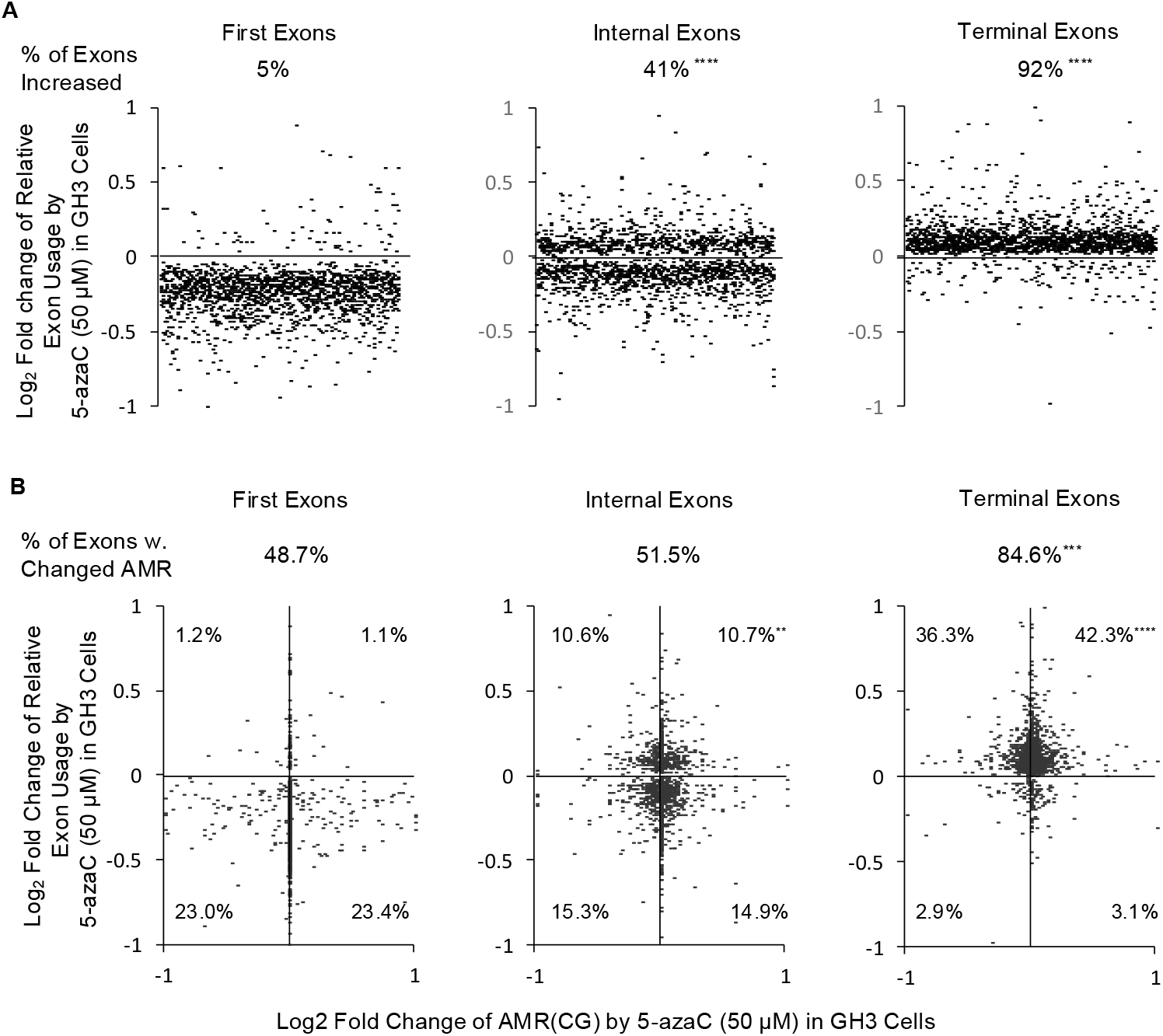
Exon usage changes of first, internal, and terminal exons following 5-azaC (50 µM) treatment in GH_3_ cells. A. Scatter plots showing log_2_ fold changes in exon usage. B. Relationship between log_2_ fold changes in exon usage and average methylation ratio (AMR). Percentages indicate exons exhibiting AMR changes upon 5-azaC treatment in each quadrant. Enrichment was assessed by hypergeometric density tests using first exons as the reference group (^**^*p* < 1.1E-37; ^***^*p* < 2.8E-120, ^****^*p* < 9.6E-162). Sample sizes were n = 1,709 first exons, 1,790 internal exons, and 1,874 terminal exons.

Moreover, we assessed the average CG methylation ratio (AMR) of exons using the samples’ available WGBS data [4]. AMR levels corroborated previous findings on exon-rank dependent methylation of expressed genes [2, 3]: most first exons were nearly unmethylated, whereas most internal and terminal exons were highly methylated in both the 5-azaC-untreated or treated cells (Fig. S1). AMR increases or decreases were distributed near evenly among exons with increased usages in each exon group; the same was observed among ones with reduced usages (Fig. 1B). However, the highest proportion of exons showing AMR changes occurred in the terminal exon group (84.6%, *p* < 2.8E-120, compared to the first exons), particularly in the first quadrant, suggesting a preferential association between AMR changes and increased terminal exon usage.

### 2. Transcriptome-wide switch from proximal to distal poly(A) sites induced by 5-azaC

We next examined exon usage in 154 and 153 genes on the (+) and (-) strands, respectively, each containing at least 10 exons/bins identified by DEXSeq analysis. Analysis of 5-azaC-induced fold changes in exon-usage revealed a shift toward increased relative usage of 3′ exons, including terminal exons, in 102 genes. For example, in the *Nfx1* gene upon 5-azaC treatment, the read ratios of the first 16 exons relative to untreated controls were mostly below 1 (Fig. 2A). In contrast, the read ratio for all downstream 3′ exons shifted to values greater than 1, indicating increased expression of a longer *Nfx1* isoform upon 5-azaC treatment. This isoform switch was independently confirmed by quantitative real-time PCR (qPCR) of RT products (Fig. 2B; p < 0.01, n = 3). The resulting longer isoform encodes an NFX1 protein containing an additional R3H domain (Fig. 2C), which binds single-stranded DNA or RNA [7], including hTERT mRNA in HPV16 E6-mediated enhancement of hTERT expression [8].

**Fig. 2.**
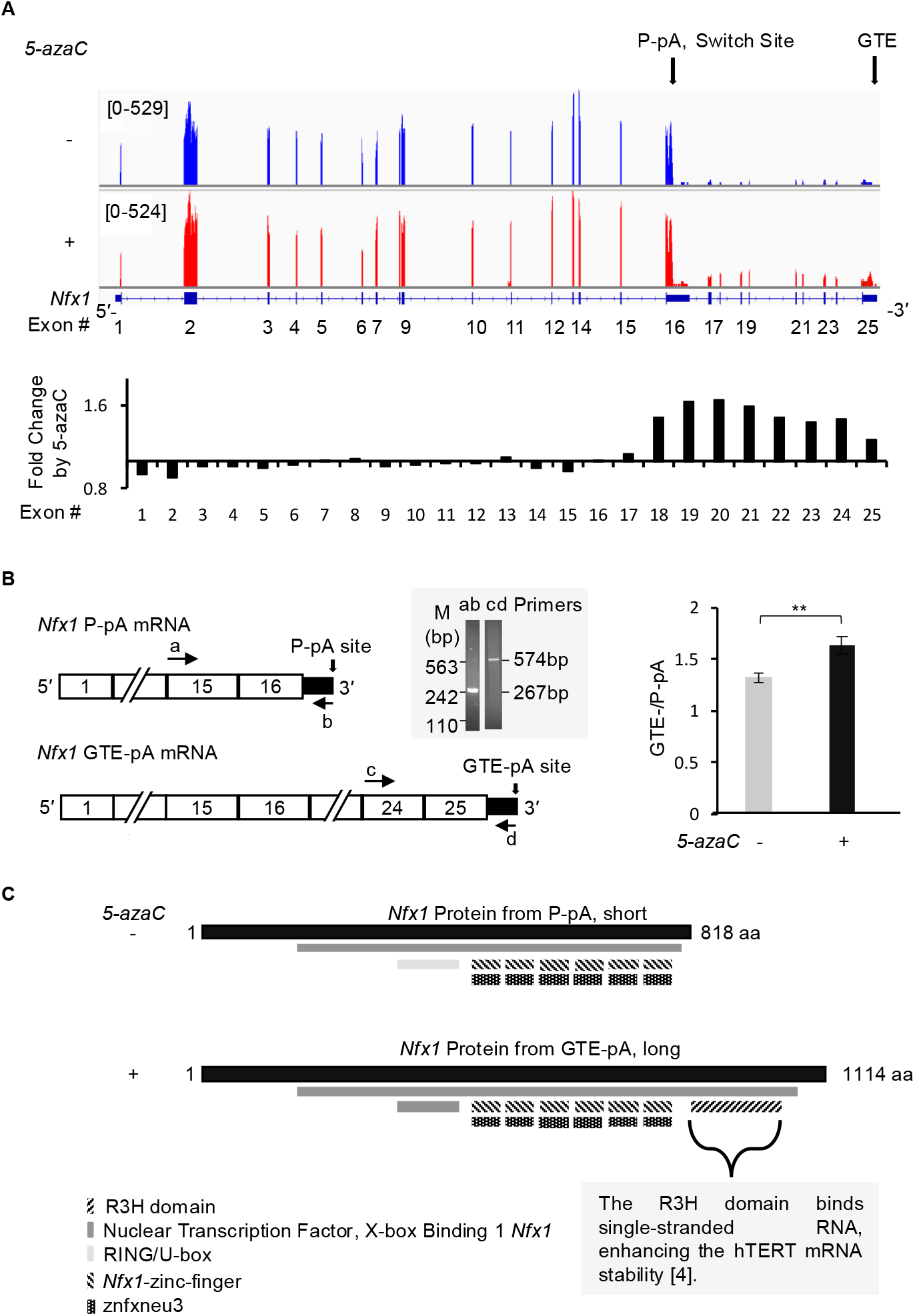
Exon rank-dependent effects of 5-azaC (50 μM) on exon usage in GH_3_ cells A. IGV view and corresponding bar plot showing the effect of 5-azaC on exon usage across the *Nfx1* gene. Bar plots of fold changes were derived from DEXSeq analysis. B. Quantitative real-time PCR (qPCR) validation of reverse-transcribed cDNA demonstrating reduced relative usage of the proximal poly(A) site and increased usage of genomic terminal exon (GTE) poly(A) sites in *Nfx1* following 5-azaC treatment (n = 3). PCR primer positions are indicated by arrowheads, and single specific products from the primers were confirmed by agarose gel electrophoresis. C. Protein domain architecture of the resulting short and long NFX1 protein isoforms, translated from transcripts ENSRNOT00000105959.2 (short) and ENSRNOT00000060692.6 (long), respectively, as annotated in the InterPro protein family database.

Similarly, the *Zmym3* gene exhibited a poly(A) site switch following 5-azaC treatment (Fig. S2A). In untreated samples, RNA-seq reads corresponding to the first six exons were approximately fivefold higher than those of the downstream 3′ exons. Upon 5-azaC treatment, expression of the 3′ exons increased to levels comparable to those of the 5′ exons, with relative usage ratios shifting from below 1.0 to above 1.0 at the junction between exons 6 and 7, indicative of a switch to the distal poly(A) site. The resulting long isoform encodes 13 additional protein domains, including DUF3504 implicated in DNA damage repair [9] (Fig. S2C).

Together, these findings suggest that 5-azaC-induced switches of alternative polyadenylation (APA) sites preferentially generate longer protein isoforms containing complete functional domain sets. These domains are among proteins with such functions as ATP binding, microtubule cytoskeleton organization, ATPase activity, and microRNAs in cancer (Table S1), as shown in the DAVID functional clustering analysis of the 102 genes.

In addition to single APA switches, a subset of genes displayed multiple switching events accompanied by alternative splicing, including exon skipping and intron retention at the switch regions. For example, intron 2 retention was detected in the *Gln1* gene following 5-azaC treatment (Fig. S3A). Notably, regions undergoing poly(A) site switching also showed pronounced changes in exon AMR. In particular, *Plekha5* and *Pafah1b1* exhibited marked AMR alterations coinciding with their switch regions (Fig. S3B, red lines in graph). Moreover, 82 of the 102 genes (80%) with single APA switches displayed significant AMR changes at the corresponding switch sites, supporting a potential mechanistic link between DNA methylation dynamics and poly(A) site selection.

### 3. Differential enrichment of poly(A) signal motifs at the proximal and terminal exon’s poly(A) sites

We next asked whether the two poly(A) sites in the 102 genes possess distinct poly(A) signal features. To address this, we analyzed consensus poly(A) signal motifs using MEME motif analysis[10], based on 200-nucleotide regions flanking the endpoints of RNA-seq read-counts visualized in IGV (Fig. 3A). We found that the consensus motif at the proximal site was relatively suboptimal compared with that at the GTE site. In addition, both sites were enriched for related but slightly divergent G-rich motifs.

**Fig. 3.**
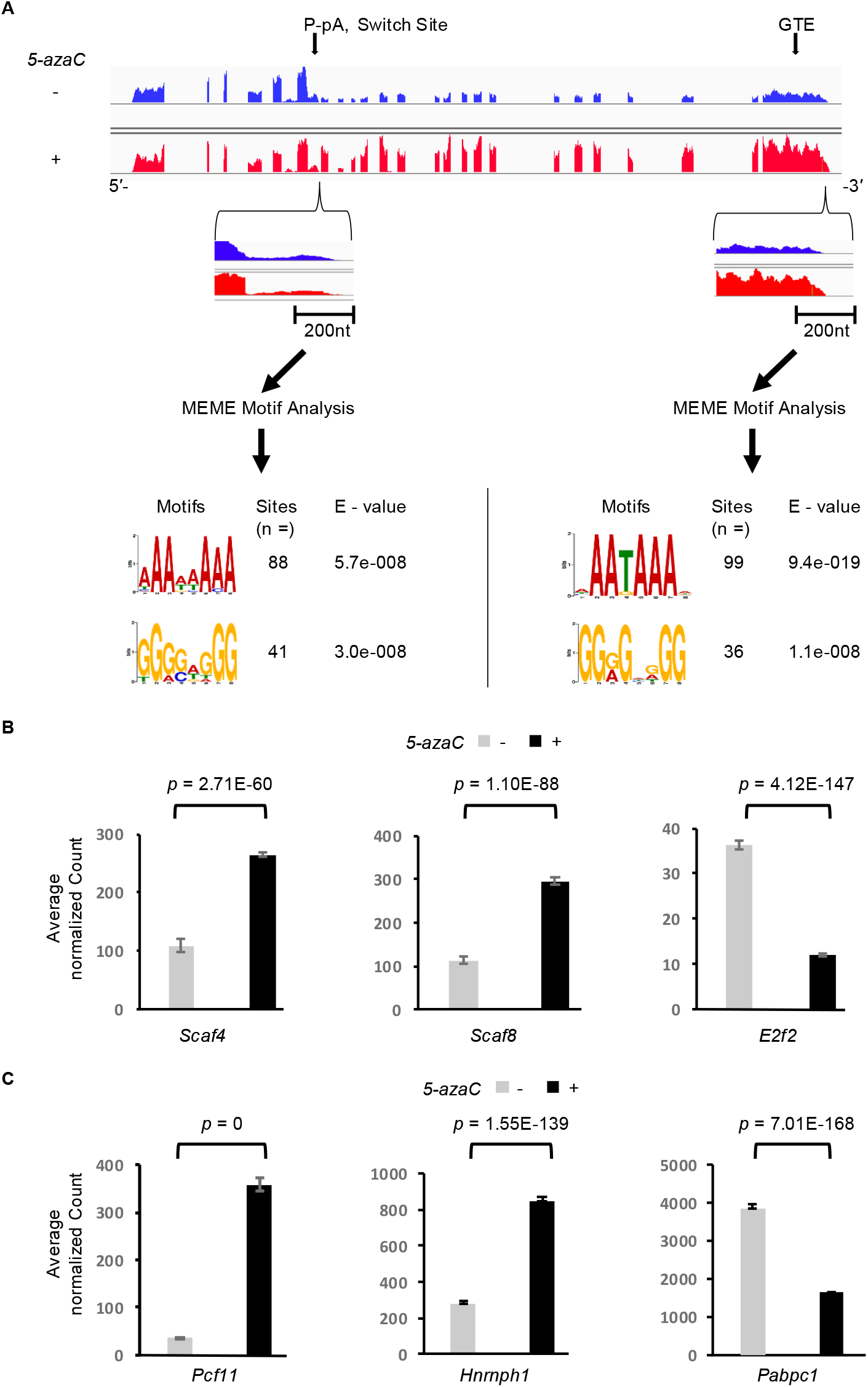
Consensus motifs of alternative termination sites and mRNA transcripts altered by 5-azaC (50 μM) in GH_3_ cells. A. Flow chart illustrating the selection of 200-nt sequences flanking read termination sites from 102 genes exhibiting a single termination-site switch for MEME motif analysis (n denotes the number of sequence sites contributing to motif construction). B. Bar plot showing average normalized counts of transcription regulatory factors (Scaf4, Scaf8, and E2f2) significantly altered by 5-azaC treatment. C. Bar plot of average normalized counts of mRNA polyadenylation and splicing factors (Pcf11, Hnrnph, and Pabpc1) significantly regulated by 5-azaC. Differential expression was determined by edgeR analysis of RNA-seq data (FDR < 0.01, 0.5 < fold change < 2, ARC > 200).

To further assess transcript termination patterns, we manually inspected RNA-seq read ends at the GTEs of 300 genes randomly selected from the 1,874 identified GTEs using IGV. Among these genes, 52% of read ends aligned precisely with the annotated gene end, 38% extended beyond, and only 10% terminated upstream the annotated end. Together, these results indicate that 5-azaC preferentially promotes full-length transcript production, primarily by shifting poly(A) site usage from proximal sites to annotated gene-end sites, and in many cases also to previously unannotated downstream poly(A) sites.

### 4. 5-azaC regulates the expression of genes that promote distal poly(A) usage

To determine whether 5-azaC-induced poly(A) site switching is accompanied by coordinated changes in relevant regulatory genes, we performed DAVID functional clustering analysis on genes significantly upregulated (fold change > 2) or downregulated (fold change < 0.5) by 5-azaC, as identified by edgeR analysis of the same RNA-seq dataset. Genes involved in transcriptional regulation emerged as a dominant functional cluster among the upregulated genes (Fig. S4), whereas genes associated with oxidative phosphorylation were prominently enriched among the downregulated genes.

Particularly, 5-azaC treatment increased the expression of multiple RNA polymerase II regulatory factors (Fig. 3B and Table S2), including Scaf4 and Scaf8, which promote distal poly(A) site usage [11]. In contrast, expression of E2f2, a factor known to favor proximal poly(A) site selection [12], was reduced. In addition, 5-azaC altered the expression of several factors implicated in alternative polyadenylation and/or splicing. Specifically, factors with reported preferences for G-rich sequences, such as Pcf11, Ssu72, Cpeb3, and Hnrnph1/h3, were upregulated, whereas Esrp2, Srsf9, Rbms2, and the A-rich–sequence–binding factor Pabpc1 were downregulated (Fig. 3C and Table S3).

Collectively, this pattern of gene regulation suggests that 5-azaC exerts a coordinated influence on transcriptional and pre-mRNA processing machinery, with the changes of at least some factors consistent with the observed global shift from proximal to distal poly(A) site usage. An apparent exception to this trend is the upregulation of Pcf11, which has been typically associated with transcript shortening and repression of distal polyadenylation sites [13], suggesting potential context-dependent or compensatory roles for this factor under 5-azaC treatment.

### 5. Similar promoting effects on terminal poly(A) sites and full-length transcripts in leukaemia cells by 5-azaC

To determine whether the ability of 5-azaC to promote full-length transcript production is specific to GH_3_ cells or also occurs in other cell types, particularly cancer cells, we analyzed raw RNA-Seq data from a study of 5-azaC-treated MOLM-13 human leukemia cells [14], deposited in the NCBI SRA (Table S4). Although a lower concentration of 5-azaC (7 µM) was used in this study [14], presence of a similar poly(A) site-switching trend would provide strong independent support for our observations in GH_3_ cells.

By using the DEXSeq analysis, we focused on genes with at least 10 exons/bins and examined exon-usage changes across first, internal, and terminal exons. Despite the reduced magnitude of the effect, likely reflecting the lower 5-azaC concentration and/or cell-type differences compared with our GH_3_ experiments (50 µM), a highly similar pattern of exon-usage shifts was observed (Fig. 4A). Specifically, fold changes for first and genomic terminal exons again shifted in opposite directions, with a highly significant enrichment of terminal exons exhibiting fold changes greater than 1.0 (*p* = 6.0E-42).

**Fig. 4.**
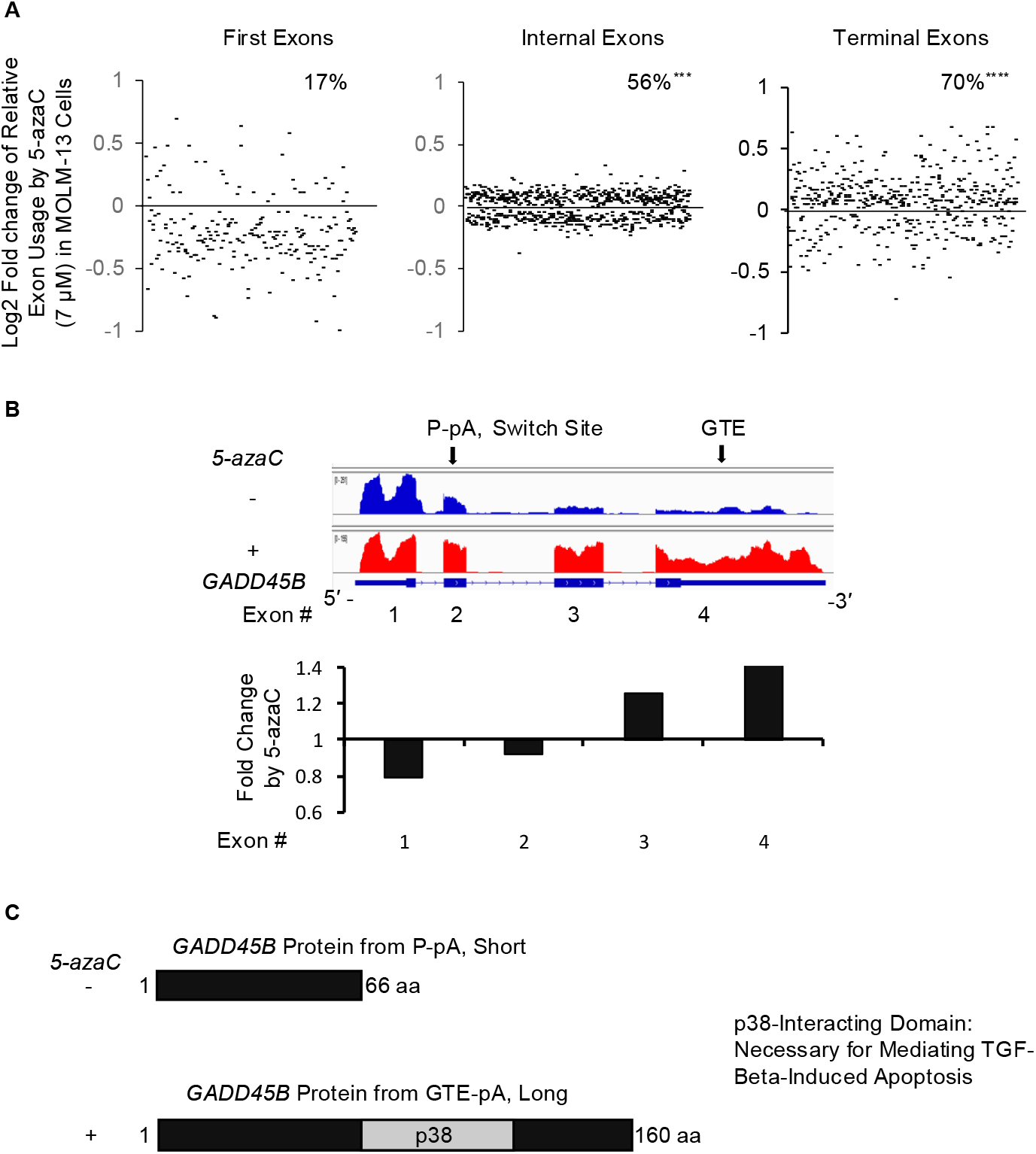
Similar 5-azaC-induced promotion of terminal polyadenylation site usage and full-length transcript production in a leukaemia cell line. A. Scatter plots showing log2 fold changes of exon usage of first, internal, and genomic terminal exons (GTEs), derived from DEXSeq analysis of RNA-Seq data from human MOLM-13 cells treated with 5-azaC (7 µM) (Zeka et al., 2023). Percentages indicate exons with fold changes > 1.0 (*p* < 0.05) in each group. Enrichment was assessed by hypergeometric density testing relative to the first-exon group (^***^*p* = 4.4E-27, ^****^*p* = 6.0E-42). Sample sizes were n = 227 first exons, 704 internal exons, and 478 terminal exons. B. Example of a poly(A) site switch to the GTE of the GADD45B tumour suppressor gene. IGV views show exon-read coverage before and after 5-azaC treatment, with corresponding bar plots of exon-usage fold changes shown below. C. Protein domains gained upon 5-azaC– enhanced terminal poly(A) site usage and full-length GADD45B transcript production. The added p38 MAP kinase-interaction domain is essential for TGF-β-induced apoptosis mediated by GADD45B.

A representative example is the tumor suppressor gene *GADD45B* (Fig. 4B-C). In untreated MOLM-13 cells, *GADD45B* was predominantly expressed as a truncated isoform, as indicated by strong RNA-seq read coverage limited to the first two exons. This shorter isoform lacks the p38-interacting domain required for mediating TGF-β-induced apoptosis [15, 16].

Following 5-azaC treatment, read coverage across exons 3 and 4 increased to levels comparable to those of the first two exons, consistent with restoration of the full-length GADD45B transcript necessary for apoptotic signaling [16].

### 6. Common changes in poly(A) factors between 5-azaC-treated cell lines

To determine whether the effects of 5-azaC on transcript levels of transcription-, polyadenylation-, and splicing-related factors are conserved between the two experimental systems, we analyzed the RNA-Seq data from MOLM-13 leukemia cells using the same edgeR analytical parameter applied to the GH_3_ cell samples. This comparison revealed that multiple transcription, polyadenylation, and splicing factors exhibited concordant regulatory trends in both cell lines, with several factors being consistently upregulated (e.g., PCF11 and CSTF3) and others downregulated (e.g., CPSF1) (Table 1). We also identified factors showing divergent regulation between the two systems. For example, CLP1, whose mouse homolog (Clp1) has been reported to promote proximal poly(A) site usage [17], was upregulated in GH_3_ but downregulated in MOLM-13 cells. Despite the differences, these results indicate that 5-azaC induces a shared regulatory group of transcriptional and RNA-processing factors that are known to influence transcript length and poly(A) site selection.

**Table 1.**
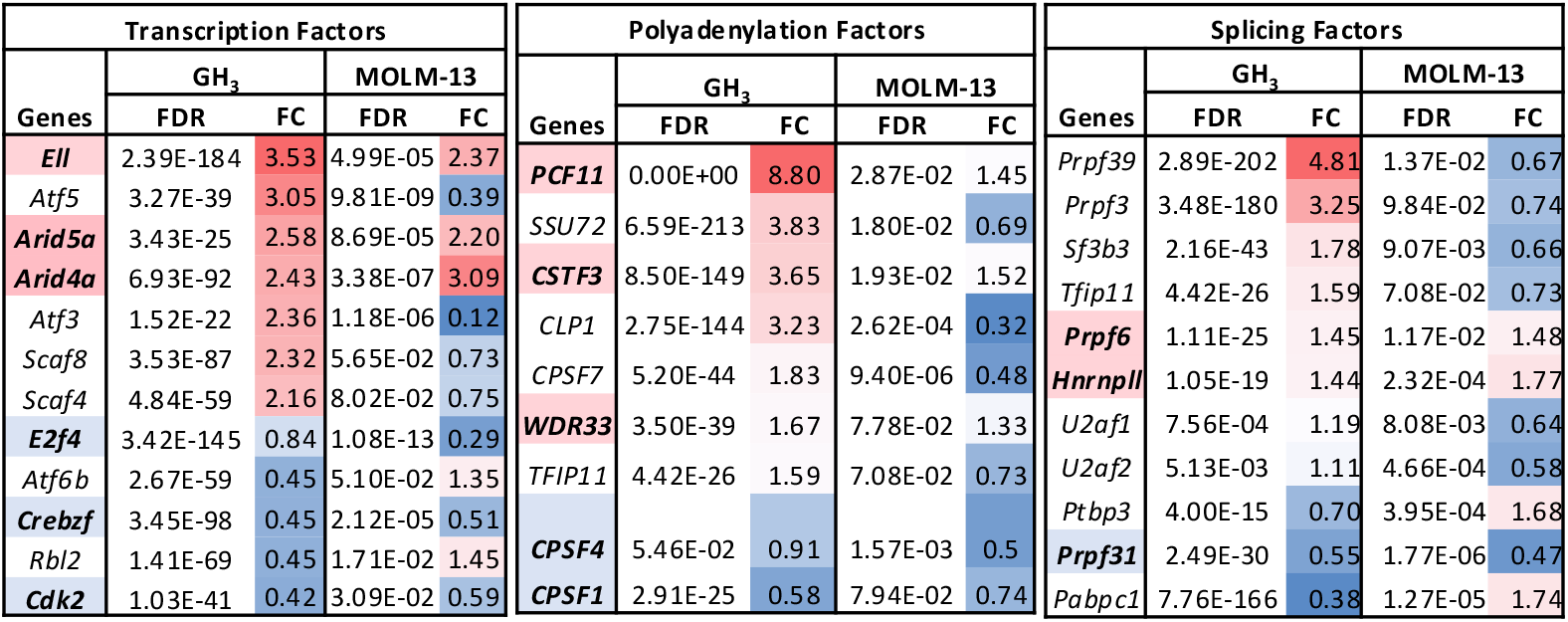
Common factors that showed changes by 5-aza C in GH_3_ (treated with 50 μM 5-aza C) and/or MOLM-13 cells (treated μM 5-aza C). Listed are changed genes from edgeR analysis of RNA-Seq data (FDR<0.1, 0.75>FC>1.25) upon 5-azaC nt. FC: fold change, FDR: false discovery rate. Genes with names in shades: RNA-Seq reads increased (pink) or reduced ue) in both cell lines.

## Discussion

In this study, we analyzed RNA-seq and whole-genome bisulfite sequencing (WGBS) data to investigate the effects of 5-azaC on exon usage and exon methylation. Our results reveal a previously unrecognized impact of 5-azaC: it extends transcript length by selectively switching poly(A) site usage from proximal sites to genomic terminal exon (GTE) sites, thereby enhancing the coordinated usage of multiple downstream 3′ exons. Although 5-azaC is known to influence alternative splicing [4], the underlying mechanisms remain poorly understood. Our data indicate a pronounced exon rank-dependent effect associated with alternative polyadenylation and a string of 3′ exons, characterized by reduced relative usage of first exons and increased usage of downstream 3′ and terminal exons (Fig. 1A). Interestingly, changes of exonic DNA methylation upon 5-azaC treatment also occurred on most of the terminal exons, and as reported [4], inversely correlate with exon usage changes at extreme levels (Fig. 1B). Together with consistent methylation changes surrounding most poly(A) switch sites (Fig. S2B), these observations imply a potential role for exon DNA methylation in poly(A) site selection.

Alternative polyadenylation is regulated by both *cis*-acting pre-mRNA motifs and *trans*- acting factors [18]. In addition to exon methylation changes (Fig. S2B), we observed distinct G-rich and consensus poly(A) motifs at proximal versus GTE sites (Fig. 3A), as well as significant alterations in poly(A) site selection factors within the transcriptional machinery. Particularly, the upregulation of Scaf4 and Scaf8 and downregulation of E2f2 following 5-azaC treatment (Fig. 3B; Table S2) are consistent with their known roles in promoting distal poly(A) usage [11]. In contrast, Pcf11, a factor typically associated with transcript shortening, is upregulated by 5-azaC in both GH_3_ and MOLM-13 cells (Fig. 3C; Table 1; Table S3)) [13].

This apparent contradiction may reflect a homeostatic response by tumor cells attempting to preserve shortened transcripts in the presence of the anti-cancer drug 5-azaC.

The unidirectional effect of 5-azaC on alternative polyadenylation, shifting poly(A) usage from proximal to terminal GTE sites, stands in sharp contrast to the transcript shortening commonly observed in cancer cells [19-21]. This distinction may represent a key component of 5-azaC’s therapeutic action, potentially restoring regulatory elements or protein functions that are lost through aberrant transcript shortening in cancer. More broadly, this newly identified effect provides a strategy to directionally extend transcript length, offering both mechanistic insight into RNA processing and a potential experimental or therapeutic avenue applicable to other existing or candidate anti-cancer drugs.

## Materials and Methods

### Cell culture

The rat GH_3_ pituitary cells were cultured as described in [4].

### 5-azaC treatment and DNA/RNA extraction

Cell cultures were treated with 5-aza-cytidine (50µM) or DMSO for 18hrs before cell collection. The cytoplasmic RNA was extracted for RNA sequencing and the corresponding nuclear DNA was extracted for WGBS analysis as described in [4, 22]. The RNA-Seq reads of 5-azaC (7µM)-treated MOLM-13 cells analysed here were obtained from PRJEB61289 deposited in the NCBI SRA database by the Pina group [14].

### Quantitative real-time PCR (qPCR)

For cDNA preparation by reverse transcription, 300ng of cytoplasmic RNA was added to a 10µl reaction, incubated at 45°C for 50 minutes and 90°C for 5 minutes. The resulting cDNA was then used to quantitate the levels of *Nfx1* proximal and GTE poly(A) site for the 5-azaC-treated and non-treated samples, using primers *Nfx1* E15F (5’-GTCCATCTGTCCTCCTACCACG-3’) and *Nfx1* E16R (5’- GTTCCAGACAGCTTACTGACG-3’) for the proximal poly(A) site and *Nfx1* E23F (5’- CATTCTTGTCATAGTGAGGAGAAG-3’) and *Nfx1* E24R (5’- GTCAACTTAACAAACAAATATGTC-3’) for the GTE poly(A) site. The samples were pre-confirmed by carrying out RT-PCR on the *Prl* gene using *Prl* -specific primers for reduced Prl expression by 5-azaC, as described previously [4]. qPCR was conducted with PowerUp™ SYBR™ Green (Applied Biosystems) on a QuantStudio 3 instrument and analysed with QuantStudio™ Design and Analysis software. Samples were amplified in triplicate and normalized to *Gh1* (growth hormone gene) using sequence-specific primers as described in [4]. Error bars represent 95% confidence intervals of the quotient of *Nfx1* GTE/proximal poly(A) site normalized to *Gh1* results.

### Bioinformatics analysis

We used IGV to visualize mapped reads [6], DAVID for functional annotation and clustering analysis of genes [23], Interpro protein family database for protein domain search [24], and MEME for motif analysis [10]. DEXSeq and edgeR analysis were carried out as in the previous studies[5, 22, 25, 26].

### Statistical tests

All samples were in triplicate. Two-tailed Student’s t-test was used for qPCR.

## Acknowledgements

This work was supported by a Discovery Grant (RGPIN-2022-05023) from the Natural Sciences and Engineering Research Council of Canada (NSERC) to J.X. S.O. is supported by a Vanier Canada Graduate Scholarship (FRN - CGV_186989). We thank Peisan Lew for assistance with qPCR.

