## Supplementary Files for "5-Aza-Cytidine Enhances Terminal Polyadenylation Site Usage for Full-Length Transcripts in Cells"

### Supplementary Figures

**Fig. S1. Methylation levels of first, internal, and terminal exons with or without 5-azaC treatment (50 µM) in $GH_3$ cells.** Scatter plots showing the average methylation ratio (AMR, CG context) of exons in $GH_3$ cells in the absence (-) or presence (+) of 5-azaC.

**Fig. S2. Exon rank-dependent effect of 5-azaC (50 µM) on the *Zmym3* gene in $GH_3$ cells.** **A**. IGV view and corresponding bar plot showing the effect of 5-azaC on exon usage across the Zmym3 gene. Bar plots were derived from DEXSeq analysis. **B**. Protein domains of the resulting short and long ZMYM3 protein isoforms, translated from transcripts ENSRNOT00000076008.3 (short) and ENSRNOT00000076004.3 (long), respectively, as annotated in the InterPro protein family database.

**Fig S3. Exon rank-dependent effects of 5-azaC (50 µM) on exon usage and DNA methylation in $GH_3$ cells.** **A**. IGV view illustrating intron retention following 5-azaC treatment, together with a bar plot showing fold changes in exon usage across the *Gln1* gene. **B**. Combined plots of exon-usage fold changes (bar charts) and net changes in average methylation ratio (AMR, CG context; line graphs) across the *Plekha5* and *Pafah1b1* genes before and after 5-azaC treatment.

**Fig S4. DAVID functional analysis of genes significantly regulated by 5-azaC (50 µM) in $GH_3$ cells.** Flow chart illustrating data selection criteria from edgeR results used for DAVID functional clustering analysis. Bar graphs display functional enrichment for genes upregulated (blue) or downregulated (brown) by 5-azaC treatment, with $-\log_{10}(p\text{-value})$ shown on the X-axis and functional categories on the Y-axis. FC: fold change, FDR: false discovery rate, ARC: average raw counts.

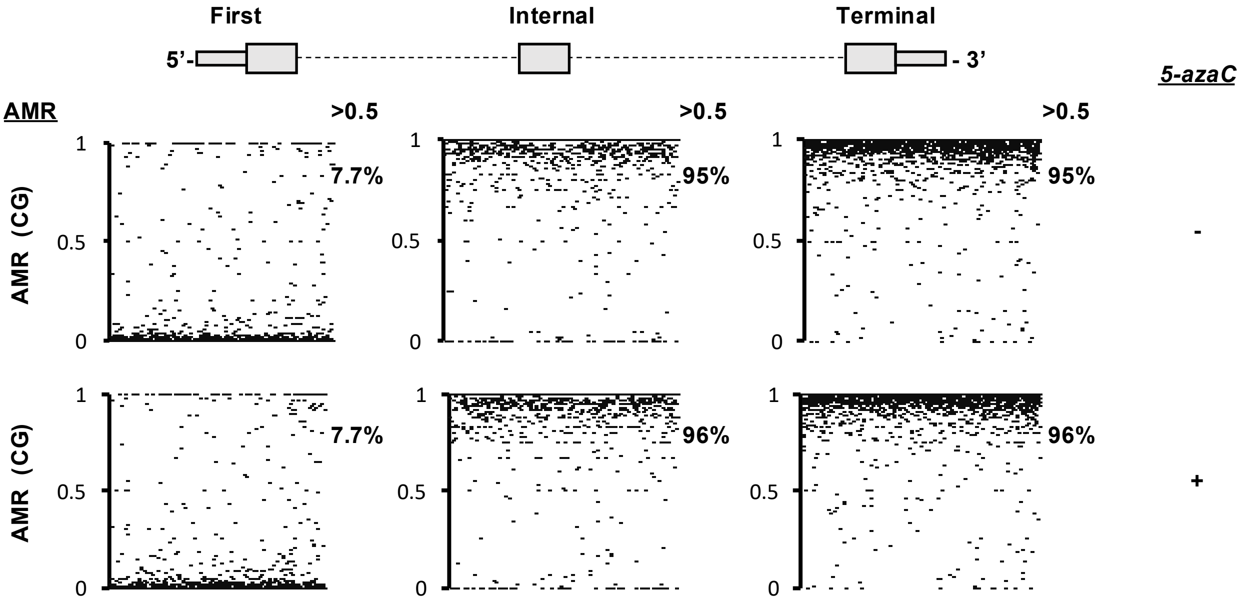
First
Internal
Terminal
5'-
- 3'
5-azaC
AMR
>0.5
>0.5
>0.5
1
0.5
0
AMR (CG)
7.7%
95%
95%
-
7.7%
96%
96%
+

**Fig.S2**

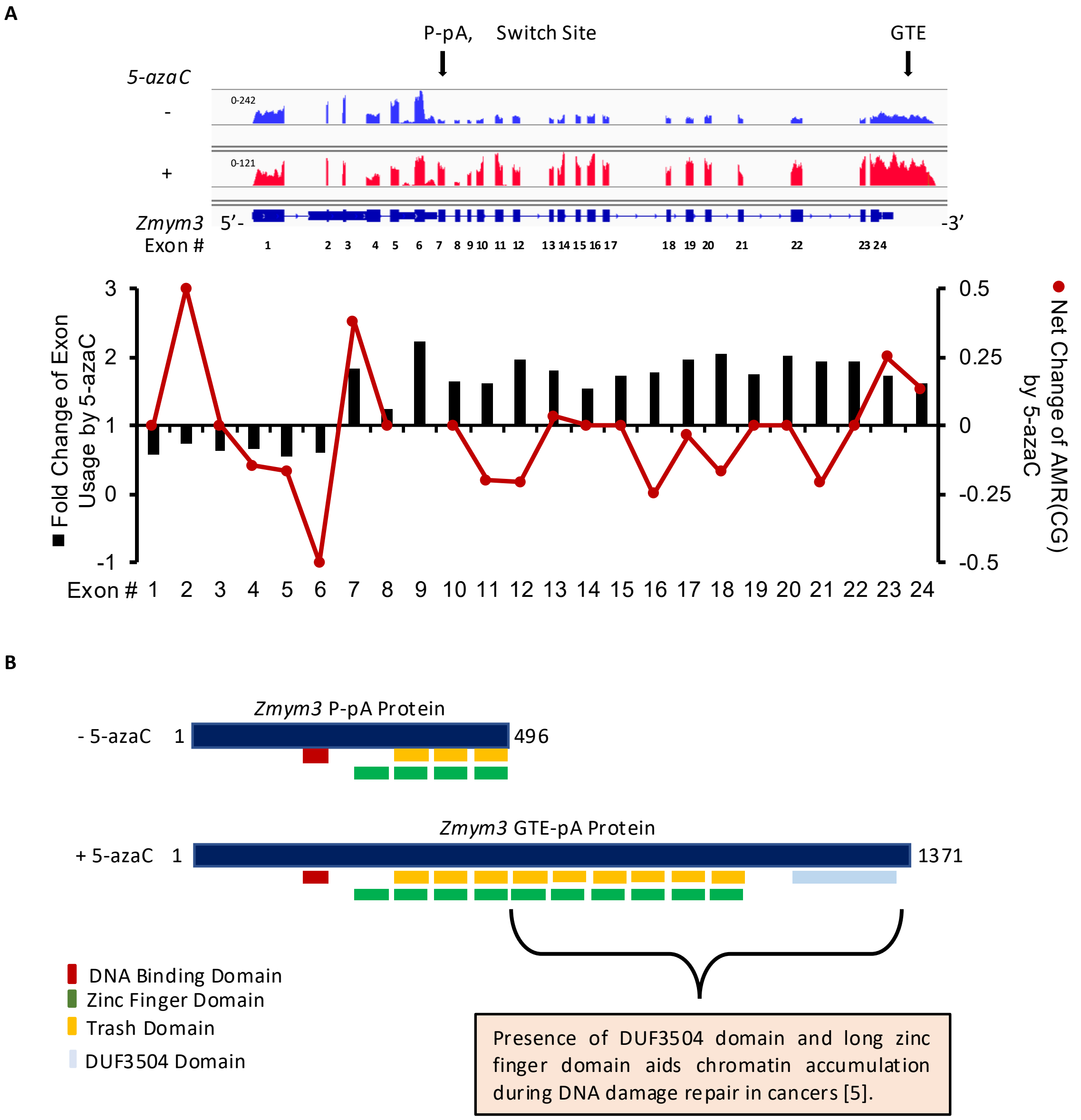
P-pA, Switch Site
GTE
5-azaC
-
+
0-242
0-121
Zmym3 5'-
-3'
Exon #
Fold Change of Exon Usage by 5-azaC
Net Change of AMR(CG) by 5-azaC
Exon # 1 2 3 4 5 6 7 8 9 10 11 12 13 14 15 16 17 18 19 20 21 22 23 24
Zmym3 P-pA Protein
- 5-azaC 1
496
Zmym3 GTE-pA Protein
+ 5-azaC 1
1371
DNA Binding Domain
Zinc Finger Domain
Trash Domain
DUF3504 Domain
Presence of DUF3504 domain and long zinc finger domain aids chromatin accumulation during DNA damage repair in cancers [5].

**A**

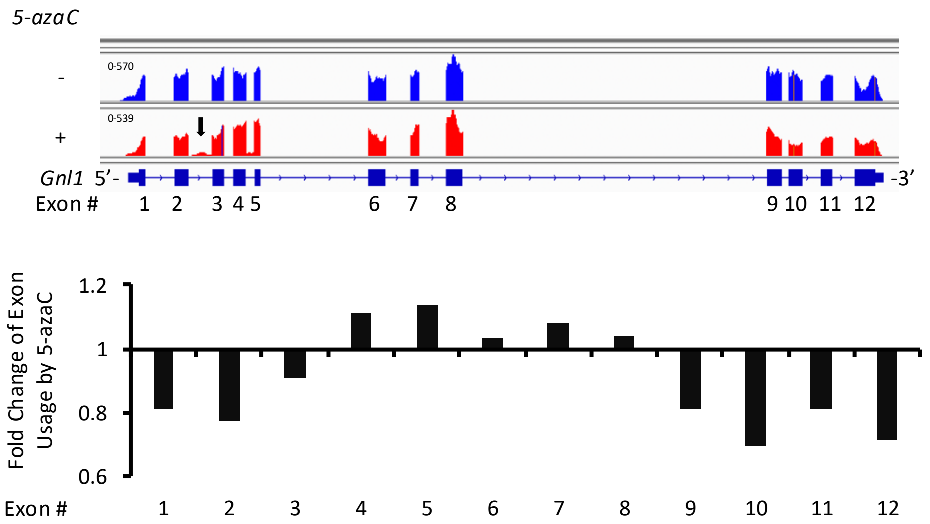
5-azaC
-
+
0-570
0-539
Gnl1 5'-
-3'
Exon #
1 2 3 4 5 6 7 8 9 10 11 12
Fold Change of Exon Usage by 5-azaC
1.2
1
0.8
0.6
Exon #
1 2 3 4 5 6 7 8 9 10 11 12

**B**

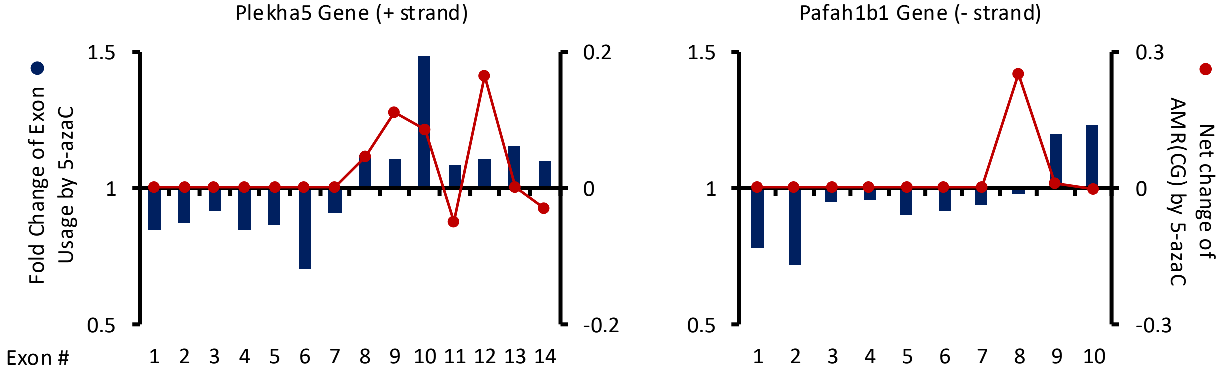
Plekha5 Gene (+ strand)
Pafah1b1 Gene (- strand)
Fold Change of Exon Usage by 5-azaC
Net change of AMR(CG) by 5-azaC
1.5
1
0.5
0.2
0
-0.2
0.3
-0.3
Exon #
1 2 3 4 5 6 7 8 9 10 11 12 13 14
1 2 3 4 5 6 7 8 9 10

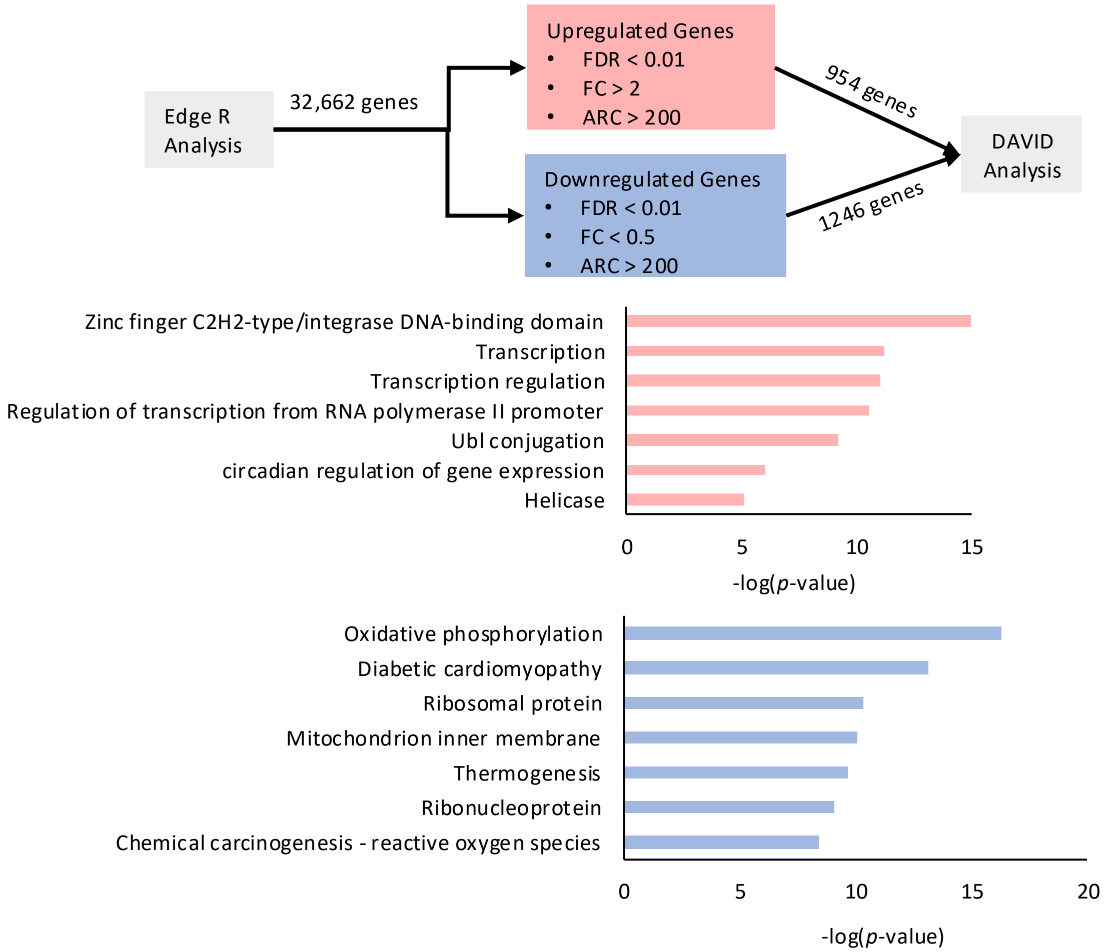
Edge R Analysis
32,662 genes
Upregulated Genes
• FDR < 0.01
• FC > 2
• ARC > 200
Downregulated Genes
• FDR < 0.01
• FC < 0.5
• ARC > 200
954 genes
1246 genes
DAVID Analysis
Zinc finger C2H2-type/integrase DNA-binding domain
Transcription
Transcription regulation
Regulation of transcription from RNA polymerase II promoter
Ubl conjugation
circadian regulation of gene expression
Helicase
0
5
10
15
-log(p-value)
Oxidative phosphorylation
Diabetic cardiomyopathy
Ribosomal protein
Mitochondrion inner membrane
Thermogenesis
Ribonucleoprotein
Chemical carcinogenesis - reactive oxygen species
0
5
10
15
20
-log(p-value)

**Table S1. DAVID functional Clustering annotation of the genes (n = 102) with single switch by 5-azaC treatment (50 µM) in GH3 cells ($p < 0.01$)**

| Categories | function | -log(p-value) | Genes |
|---|---|---|---|
| **GOTERM_MF_DIRECT** | ATP binding | 4.58502665 | *Abca3, Abcb1a, Clk2, Ddx54, Iqcd, Sik3, Shprh, Smarca2, Acsl6, Apaf1, Chd1 ,Chd4, Cars, Ppip5k1, Ern1, Egfr, Hipk1, Ikbkb, Kif1b, Kif21b, Magi1, Mark1, Myo5c, Myh14, Pgd, Gart, Ptk2, Smchd1, Trio* |
| **GOTERM_MF_DIRECT** | chromatin binding | 4.39794001 | *Crebbp, Smarca2, App, Ankrd17, Rere, Chd1, Cux1, Ezh1, Egfr, Parg, Tnrc18, Trim66, Ubtf* |
| **GOTERM_CC_DIRECT** | neuron projection | 3.88605665 | *Shank1, Shank3, App, Bptf, Cux1, Cyfip2, Gabbr1, Grin1, Kif1b, Nrcam, Pafah1b1, Tenm3* |
| **GOTERM_MF_DIRECT** | microtubule binding | 3.21467016 | *Abca3, Abcb1a, Ddx54, Iqcd, Smarca2, Chd1, Chd4, Kif1b, Kif21b, Smchd1* |
| **GOTERM_BP_DIRECT** | positive regulation of excitatory postsynaptic potential | 3.000000 | *Shank1, Shank3, App, Grin1* |
| **KEGG_PATHWAY** | MicroRNAs in cancer | 2.537602 | *Apc2, Abcb1a, Cd44, Crebbp, Egfr, Ikbkb, Slc7a1* |
| **KEGG_PATHWAY** | Human papillomavirus infection | 2.14874165 | *Apc2, Atp6v0a2, Crebbp, Chd4, Egfr, Ikbkb, Lama4, Magi1, Nfx1, Ptk2* |

**Table S2. DAVID functional Clustering annotation of the genes (n = 43) with single switch by 5-azaC treatment (7 μM) in MOLM-13 cells (p<0.01)**

| Categories | function | -log(p-value) | Genes |
|---|---|---|---|
| **GOTERM_BP_DIRECT** | cytoplasmic translation | 27.5850267 | *RPL10, RPL13, RPL26, RPL36, RPL37A, RPL41, RPL8, RPS17, RPS2, RPS24, RPS3, RPS3A, RPS5, RPS8* |
| **KEGG_PATHWAY** | Coronavirus disease - COVID-19 | 18.69897 | *RPL41, RPL8, RPS17, RPS2, RPS24, RPS3, RPS3A, RPS5, RPS8* |
| **GOTERM_MF_DIRECT** | RNA binding | 12.853872 | *FAU, ALDOA, EIF4A1, MATR3, RNASET2, RPL10, RPL13, RPL26, RPL36, RPL37A, RPL41, RPL8, RPS17, RPS2, RPS24, RPS3, RPS3A, RPS5, RPS8, SRRM2, ZFR* |
| **GOTERM_CC_DIRECT** | focal adhesion | 7.15490196 | *ACTG1, RPL37A, RPL8, RPS17, RPS2, RPS3, RPS3A, RPS5, RPS8, RPLP2* |
| **UP_KW_PTM** | Acetylation | 4.52287875 | *NDRG1, NPLOC4, ACTG1, ALDOA, CSRNP2, EIF4A1, MATR3, MAD1L1, RPS8, RPLP2, SRRM2, SLC25A3, ZFR* |
| **GOTERM_MF_DIRECT** | mRNA binding | 2.88605665 | *EIF4A1, RPS2, RPS3, RPS5, SRRM2* |
| **UP_KW_BIOLOGICAL_PROCESS** | Translation regulation | 1.88605665 | *MKNK2, RPL10, RPS3* |

**Table S3. Polyadenylation and alternative splicing factors that were significantly regulated by 5-azaC in $GH_3$ cells (treated with 50 μM 5-azaC)**. Table showing the gene name, gene ontology, FDR, and heatmap of fold change of polyadenylation and alternative splicing genes from edgeR analysis of RNA-Seq data that were significantly changed (FDR<0.01, 0.5<FC>2, ARC > 200) by 5-azaC treatment. (FC – fold change, FDR – false discovery rate, ARC – average raw counts).

| Gene Name | Gene Ontology | FDR | |
|---|---|---|---|
| *Pcf11* | mRNA polyadenylation, cleavage and processing | 0 | **8.8** |
| *Hnrnph3* | Regulation of RNA splicing | 2.24E-99 | |
| *Ssu72* | mRNA Polyadenylation | 6.59E-213 | |
| *Cstf3* | mRNA 3'-end processing | 8.50E-149 | |
| *Clp1* | mRNA 3'-end processing and polyadenylation | 2.75E-144 | |
| *Ptbp2* | mRNA splice site selection and processing | 5.28E-85 | |
| *Hnrnph1* | mRNA processing | 1.15E-137 | |
| *Cpeb3* | Positive regulation of mRNA polyadenylation | 6.31E-56 | |
| *Fip1l1* | pre-mRNA cleavage required for polyadenylation | 6.67E-50 | |
| *Cstf2t* | pre-mRNA cleavage required for polyadenylation | 7.70E-78 | |
| *Sf3a1* | mRNA 3'-splice site recognition | 4.34E-80 | |
| *Pabpc4-ps1* | RNA processing | 3.20E-17 | |
| *Mbnl1* | Regulation of alternative mRNA splicing, via spliceosome | 2.77E-61 | |
| *Srsf9* | mRNA splice site selection | 1.04E-99 | |
| *Rbms2* | RNA processing | 6.81E-62 | |
| *Pabpc1* | positive regulation of nuclear-transcribed mRNA poly(A) tail shortening | 7.76E-166 | |
| *Hnrnpa1* | Regulation of alternative mRNA splicing, via spliceosome | 3.19E-153 | |
| *Esrp2* | Regulation of RNA splicing | 4.48E-100 | **0.35** |

**Fold change of gene expression by 5-azaC Treatment**
